# Benefits of Deep Learning Classification of Continuous Noninvasive Brain-Computer Interface Control

**DOI:** 10.1101/2020.09.09.289462

**Authors:** James R. Stieger, Stephen A. Engel, Daniel Suma, Bin He

## Abstract

Noninvasive brain-computer interfaces (BCIs) assist paralyzed patients by providing access to the world without requiring surgical intervention. While the performance of noninvasive BCI is hindered by long training times and variable user proficiency, it may be improved by deep learning methods, such as convolutional neural networks (CovNets). Prior work has suggested that the application of deep learning to EEG signals collected over the motor cortex during motor imagery based BCI increases classification accuracy in standard sensorimotor rhythm (SMR) BCI datasets. It remains to be seen whether these improvements can generalize to practical scenarios such as continuous control tasks (as opposed to prior work reporting one classification per trial), or whether valuable information remains latent outside of the motor cortex (as no prior work has compared full scalp coverage to motor only electrode montages). Here we report that deep learning methods significantly increase offline classification accuracy on an independent, large, and longitudinal online motor imagery BCI dataset with up to 4-classes and continuous 2-dimensional feedback. Improvements in classification accuracy were found to negatively correlate with initial online BCI performance, suggesting deep learning methods preferentially benefit BCI participants who need it most. The CovNets also significantly increased the information transfer rate (ITR) of the BCI system: They produced a two-fold increase in ITR without loss in classification accuracy when comparing CovNet models trained with full scalp EEG coverage to the traditional motor cortex specific decoding. Our results suggest that a variety of neural biomarkers useful for BCI, including those outside the motor cortex, can be detected through deep learning methods.

## 1. Introduction^1^

Those afflicted by paralysis resulting from stroke, trauma, and other neuromuscular disorders suffer from reduced quality of life (French et al., 2010). The ability to noninvasively decode user intent in real time enables brain-computer interfaces (BCIs) to provide this population with alternative routes of communication and action (He et al, 2020). Decoding requires the recording of neural signals, often through electroencephalography (EEG) due to its low cost, portability, and non-invasive nature. Various types of EEG based BCIs have been developed to detect different neural biomarkers, and each has their own benefits and drawbacks. BCIs based on exogenous stimulation—for instance, the P300 response (Mak et al., 2011) and steady state evoked potentials (Chen et al., 2015)—present the user with a display and use the visual system’s response to control a device. While these systems often work well, they require a visual display with pre-programmed options, making them unsuitable for practical continuous control. BCIs based on the endogenous modulation of brain rhythms—in response to motor imagery (Wolpaw and McFarland, 2004), shifts in visual attention (Tonin et al., 2013; Treder et al., 2011; van Gerven and Jensen, 2009), and manipulations of high level attention (Grosse-Wentrup and Schölkopf, 2014)—are intuitive and provide continuous control, but may be more challenging to learn. Finally, some BCIs have been proposed which combine multiple biomarkers such as responses to both motor imagery and overt shifts in attention (Li et al., 2010; Meng et al., 2018).

One well-studied BCI method attempts to decipher motor imagery by recording sensorimotor rhythms (SMRs), which predictably change in response to real and imagined movements (Pfurtscheller and Neuper, 2001; Yuan et al., 2009). One of the most prominent SMRs is the mu rhythm, a characteristic biomarker of the motor cortex at rest, visible as oscillations in the alpha band over midline structures. When moving, or thinking about moving, the motor cortex is engaged, leading to reductions in the mu rhythm contralateral to the moved, or imagined, body part; this phenomena is called event-related desynchronization (ERD). The reliability of ERD and its converse, event-related synchronization (ERS—increases in alpha power over the motor cortex during rest), can be used to control a BCI (Pfurtscheller and Lopes Da Silva, 1999; Pfurtscheller and Neuper, 2001). The intuitive and continuous nature of SMR-based BCIs enables the extension of the user through the control of virtual objects, drones, wheelchairs, and robotic arms (Doud et al., 2011; Edelman et al., 2019; Huang et al., 2012; Lafleur et al., 2013; Meng et al., 2016; Rebsamen et al., 2010; Wolpaw and McFarland, 2004), and has further been shown to improve post-stroke outcomes when combined with physical therapy (Frolov et al., 2017).

Unfortunately, to realize the full potential of SMR based BCIs, a deeper understanding of the nature of brain activity and user intent associated with specific BCI tasks is required. This deficit is demonstrated by long training times and variable user control quality, which prevent the widespread clinical adoption of BCIs (Leeb et al., 2013). At least 20% of users are unable to control SMR BCIs proficiently, even after extensive training. Furthermore, performance levels often fluctuate, making the safe use of robotic end effectors for daily activities difficult (Guger et al., 2003). Why some users are unable to control the device and how to remedy this situation is currently unknown, although some psychological and electrophysiological factors correlate with initial BCI skill (such as higher resting alpha power) (Ahn et al., 2013b; Blankertz et al., 2010; Hammer et al., 2011).

Further improvement in SMR based BCIs may be realized through the application of deep learning with neural networks. The promise of deep learning lies in its capacity for learning complex nonlinear patterns from the raw data (Craik et al., 2019). Many recent studies have shown that neural networks are able to classify user intent from brain data with excellent accuracy (Craik et al., 2019; Lu et al., 2017; Sakhavi et al., 2018; Schirrmeister et al., 2017) as well as improve the information transfer rate (ITR) of BCI systems (Nagel and Spüler, 2019). However, many of these studies rely on the same public BCI competition datasets, or small data sets using simple trial structures, and therefore may not generalize well outside the datasets they were tested on (Jayaram and Barachant, 2018). Most of these studies utilize the BCI Competition IV datasets 2a and 2b, which are comparatively small (9 subjects, 2-5 sessions) and simple (2a—4 class, no online feedback, 2b—2 class with online feedback, but only 3 motor electrodes) (Tangermann et al., 2012).

Finally, most deep learning BCI classification studies have relied on a uniform trial structure and utilized the entire trial for classifications, which is not suitable for continuous BCI control (e.g., classifying a 2s trial of motor imagery as “left” rather than continually updating a cursor position in response to a prediction signal) (Wolpaw and McFarland, 2004). While online deep-learning BCI systems have been recently developed, these systems have yet to demonstrate continuous feedback control. (Jeong et al., 2020; Tayeb et al., 2019). For example in Tayeb et al., 20 subjects performed 2 class motor imagery (left hand, right hand) and at the end of the trial, a robotic arm was moved in the direction classified by the system. Jeong et al. developed a system closer to continuous control using a moving window approach where 15 subjects performed 6 class motor imagery (imagine the right hand moving left, right, up, down, forward, and backward). Sliding windows of 4s were updated every 0.5s, and classifier output values were averaged over 3 successive windows and the direction with the highest average probability over these windows was selected as an output command for a robotic arm. Unfortunately knowing a desired direction of movement is not the same as fluidly moving an external device through space (Edelman et al., 2019). Therefore, rigorous validation of previous methods and methodological extensions suitable for online continuous control are needed to aid the development of online BCI systems using deep learning classification.

This work addresses two questions that need to be answered before the utility of deep-learning can be fully established: 1) Does deep learning generally improve BCI classification, and if so, how? 2) How do we bridge the gap between offline and single decision BCI classification studies and online continuous control BCIs based on deep learning classification? We recently collected, to our knowledge, the largest motor imagery BCI dataset ever recorded (in terms of total recording time; e.g., roughly 950% larger than the BCI competition datasets 2a and 2b combined) which includes 76 subjects and up to 11 sessions of online, continuous BCI control (Stieger et al., 2020). In each session, participants performed 300 trials of up to 4 class motor imagery in center-out cursor tasks with online feedback. Factors, besides size, that separate our dataset from those examined previously include: 1) the recording of high density (64 channel) EEG, 2) varying trial structure (i.e., trials vary in length between 0.5s and 6s rather than a uniform duration of 2 or 4s), and 3) continuously updated online feedback in 1D and 2D. Validating deep learning methods on this dataset presents a unique opportunity to study how neural networks classify individuals’ brain data from the population at large, under challenging conditions (such as 4 class, center-out, continuous control with varying trial structure), and additionally represents the first assessment of how neural net classification accuracy changes based on the inclusion of all electrodes provided by high density EEG (Figure 1).

**Figure 1.**
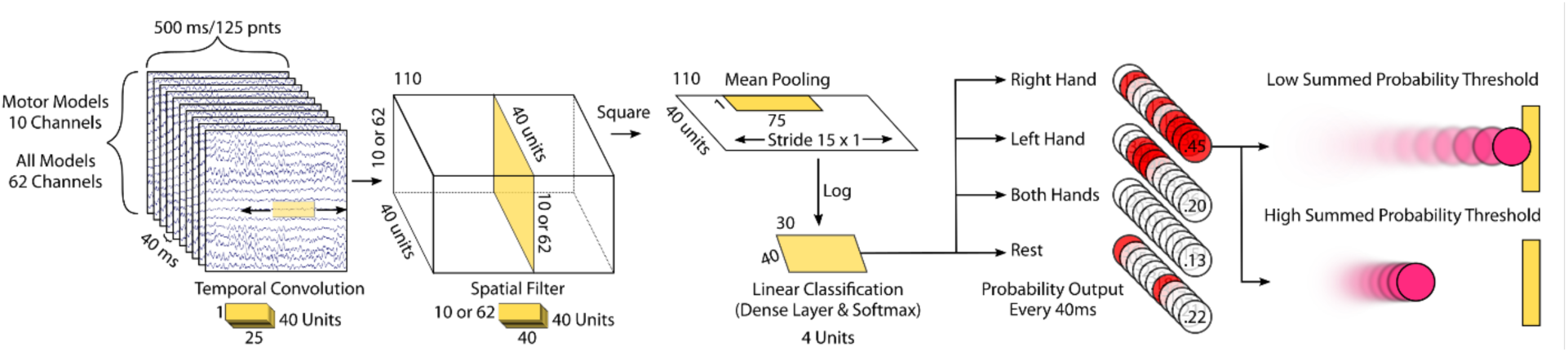
Continuous Deep Learning Classification of BCI Data. Sliding windows of EEG data recorded during online BCI tasks were used to train the shallow CovNet architecture reported in Schirrmeister et al., 2017. These windows were 500ms long and were shifted every 40ms. Two types of models were trained based on the data provided. “Motor Models” were trained using the same motor cortex electrode montage as the online BCI experiments. “All Models” were trained using all available elctrodes. In successive steps, the shallow CovNet architecture performs a temporal convolution, spatial filter, squaring nonlinearity, mean pooling, log transform, and linear classification using a dense layer and softmax transform. During testing, the trained models provide the estimated probability of class membership for each window. In a simulated cursor control environment, the class with the highest estimated probability (red circles) was used to move a virtual cursor in the direction of, and in proportion to, this highest estimated probability. The functional mapping between neural network predictions and control systems was explored by varying the summed probability threshold (the sum of the neural network outputs over time) needed for trial classification. Low probability thresholds emulated faster cursor control and higher thresholds emulated slower cursor control (see text for further details).

## 2. Materials and Methods

### 2.1. Dataset

During the BCI tasks, users attempted to move a virtual cursor outwards from the center of the screen to one of four targets placed at the cardinal positions (Top, Bottom, Left and Right). Participants were instructed to imagine their left (right) hand opening and closing to move the cursor left (right), imagine both hands opening and closing to move the cursor up and finally, rest, to move the cursor down. In separate blocks of trials, participants drove the cursor toward a target that required left/right (LR) movement only, up/down (UD) only, and combined 2D movement (2D). BCI experiments consisted of 3 runs of the LR, UD, and 2D tasks (each with 25 trials), a short break, and an additional block of 3 runs of each task. Runs were broken into individual trials which included pre and post inter-trial intervals (2 s each), a cuing period where only the target was presented (2 s), and up to 6s where the participant was provided feedback and attempted to control the cursor (0.04-6 s). Only the feedback portion of trials was used for classification. Trials longer than 6s were aborted with the closest target to the current cursor position selected as the intended target.

BCI experiments were conducted using BCI2000, a general-purpose software platform for BCI research (Schalk et al., 2004). The control signal was extracted as different combinations of the autoregressive (AR) spectral amplitudes of the small Laplacian filtered electrodes C3 and C4 in a 3 Hz bin surrounding 12 Hz, normalized to zero mean and unit variance. The magnitude of the cursor movement was determined by the normalized AR amplitude difference in a 160ms window, updated every 40ms. Horizontal motion was controlled by lateralized alpha power (C4 - C3) and vertical motion was controlled by up and down regulating total alpha power (C4 + C3).

In total, 76 participants finished all experimental requirements. The first group of participants (n = 21) completed 7 BCI sessions, however results indicated learning was still taking place; thus, the experiment was extended. Four subjects were excluded from the analysis due to non-compliance with the task demands and one was excluded due to experimenter error. Further, to evaluate the impact of deep learning classification on those not immediately proficient in standard BCI control, analysis focused on those that did not demonstrate ceiling performance in the baseline BCI assessment (accuracy above 90% in 1D control, measured by percent valid correct—hits/(hits+misses) i.e., excluding aborts). Therefore, the results presented here describe data collected from 62 participants. The study was approved by the institutional review boards of Carnegie Mellon University and the University of Minnesota, and all participants provided written informed consent.

Minimal preprocessing was applied to the dataset, and could, in theory, be implemented online (Daly et al., 2015; Jafarifarmand and Badamchizadeh, 2019). EEG data were bandpass filtered between 1 and 100 Hz, and then down-sampled from 1kHZ to 250 Hz. Noisy channels, identified through visual inspection, were replaced by local weighted averages interpolated through spherical splines (Delorme and Makeig, 2004; Ferree, 2006). The data were re-referenced to a common average. Ocular artifacts were removed using independent component analysis (ICA) and a template matching procedure. Trials shorter than the minimum window length were removed (Stieger et al, 2020).

### 2.2. CovNet Models

In order to emulate BCI experiments in which deep learning could be used in an online setting after calibration, one model was trained for each session using the first two runs in a given block and the last run in each block for testing (i.e., 2 training splits per session). Electrode-wise exponential moving standardization with a decay factor of 0.999 was used to compute exponential moving means and variances for each channel and standardize the continuous data (Schirrmeister et al., 2017). Convolutional neural network (CovNet) models were trained using the brain_decode package (version 0.4.85) in python, specifically using their shallow CovNet architecture (Schirrmeister et al., 2017). Briefly, in this architecture, the first two layers perform a temporal convolution and a spatial filter followed by a squaring nonlinearity, mean pooling layer, and a logarithmic activation function (Figure 1). Due to the variable trial structure and an attempt to reproduce the online classification, we used the recommended default time window length from BCI2000 of 500ms, shifted in increments of 40ms such that one estimation of the probability of class membership was made every 40ms. To compare this output to the classification of the online system in the 1D case, non-valid classes were removed from the CovNet predictions after classification (e.g., in the LR task the classification probability for up/down trials was set to zero). Two different models were trained for each session. The first model (Motor-Models—10 electrodes) used only the electrodes over the motor cortex (C3/C4 each surrounded by the 4 electrodes that formed the small Laplacian filter for online control), while the second included all available electrodes for training and testing (All-Models—62 electrodes). Models were trained using the Extreme Science and Engineering Discovery Environment (XSEDE), which is supported by National Science Foundation grant number ACI-1548562. Specifically, it used the Bridges system, which is supported by NSF award number ACI-1445606, at the Pittsburgh Supercomputing Center (PSC) (Nystrom et al., 2015; Towns et al., 2014).

### 2.3. Information Transfer Rate

ITR is a useful measure in comparing BCI systems because it combines both accuracy and speed of use into one metric, and can be defined (Wolpaw et al., 2002) as bits/trial with the following equation:

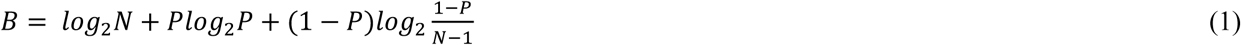

Where *B* is the ITR in bit rate (bits/symbol), *N* is the number of possible target choices, and *P* is the probability that the desired choice will be selected, otherwise called the classification accuracy. Typically, ITR *B*_*t*_ is reported units of bits/min, and calculated in (2) with *T* (seconds/symbol) being the average trial length, or time needed to convey each symbol.

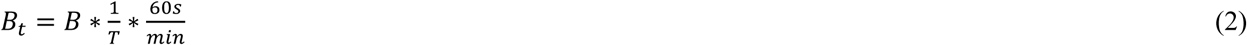

### 2.4. Electrophysiological Analysis

Feature importance maps were calculated with the input-perturbation network-prediction correlation method (Schirrmeister et al., 2017). These maps were created to investigate the causal effect of changes in EEG power on the trained CovNet models. Initially, all 500ms windows were transformed to the frequency domain by a Fourier transform. Then the power amplitudes were randomly perturbed by adding Gaussian noise (with mean 0 and variance 1) to their original values while leaving phase unperturbed. These trials were then transformed back to the time domain via an inverse Fourier transform. Predictions were made by the trained CovNet models before and after the perturbations. This procedure was repeated 400 times, then the change in input amplitudes (the added noise) and the change in CovNet predictions was examined in a correlation analysis. Feature importance maps were calculated only for the final 3 sessions when learning was assumed to have plateaued, then averaged for each subject. A cluster-based permutation test statistic was used to assess group differences in feature importance maps for each target condition (Maris and Oostenveld, 2007; Oostenveld et al., 2011).

In order to assess the electrophysiological response to BCI feedback, epochs were collected around time points of the greatest horizontal cursor movement. A time course of the cursor velocity—the derivative of the cursor positions provided by BCI2000—was first extracted and the positive peaks of this signal were identified (i.e., the local maxima of cursor velocity toward the intended target). These peak velocity measures were ordered in terms of magnitude and the 100 greatest peaks from each run (600 per session) were selected as time points of interest. Epochs were created by collecting the data in a 1.5s window (−0.5s—1s) surrounding these peaks, and neural responses were averaged across these epochs in each session.

Complex Morlet wavelet convolution was used to estimate the power spectrum of the EEG signals. A family of wavelets was created with 30 frequencies log spaced between 4 and 60 Hz with cycles increasing from 3 cycles at 4 Hz to 10 cycles at 60 Hz. The alpha band power was extracted by averaging a 3 Hz bin centered at 12 Hz.

### 2.5. Statistics

Most data displayed significant deviations from normality and were therefore fit with a rank based analysis of linear models using the Rfit package (0.23.0) as a robust alternative to least squares (RAOV) (Kloke and McKean, 2012). Metrics were fit in 2×2 designs with levels of classification type (3 Levels: Online, Motor-Models, and All-Models, or 2 Levels: Motor-Models, and All-Models) vs. task type (3 Levels: LR, UD, 2D). Significant findings were followed by independent post-hoc tests to aid inference. All post-hoc inferences were two-sided at a Holm-Bonferroni corrected alpha level of 0.05 (0.025 per side).

### 2.6. Data and code availability statement

The code and data used in this work will be made available upon reasonable request and may not be used for commercial purposes.

## 3. Results

### 3.1. CovNet Model Predictions Outperform Online Classification Accuracy

Deep learning with CovNets was found to significantly improve classification accuracy across all three tasks. Initially, CovNet predictions were averaged across all windows of a given trial and the class of the highest average probability was chosen as the classification for a given trial. Figure 2A displays the classification accuracy of the online experiments, motor electrode only trained CovNet models (Motor-Models), and CovNet models using all electrodes (All-Models) throughout participant training. By examining performance in each task across participant training sessions, two facts become apparent. First, this improvement in BCI classification accuracy is maintained across all tasks and all sessions, with a slight benefit given to models using all electrodes. Second, as participants learn to produce better motor imagery (and brain signals), the CovNets are able to discern this improvement, which is evinced by increases in classification accuracy over time.

**Figure 2.**
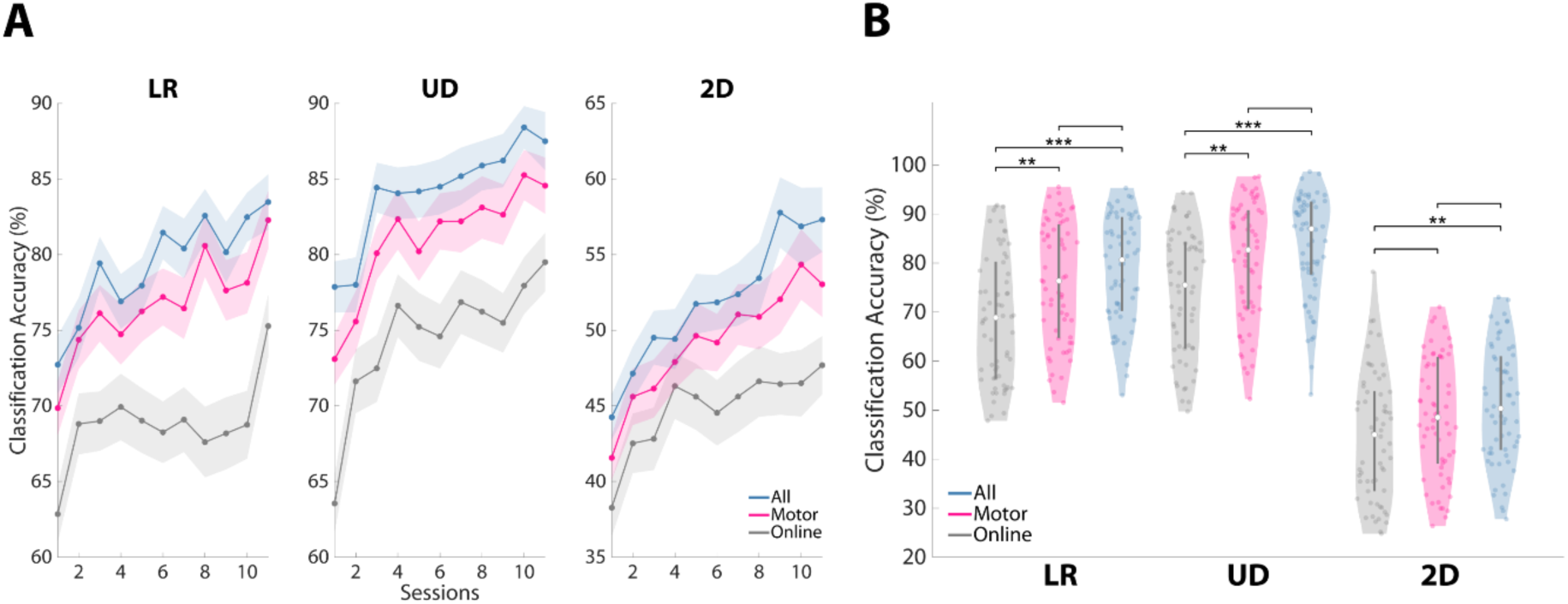
CovNet Model Predictions Outperform Online Classification Accuracy. **A**, The use of CovNet Models was found to increase classification accuracy in all tasks examined throughout BCI training. CovNet models with all electrodes (blue) displayed the best classification accuracy in each individual session, followed by motor electrode only CovNet models (pink) and online classification accuracy with BCI2000 (grey). **B**, Significant improvements in the average BCI classification accuracy were found for all tasks when using full scalp EEG coverage (All-Models: LR (WRS, P=0.00002), UD (WRS, P=0.00002), and 2D (WRS, P=0.006)), however significant improvements were only seen in the 1D tasks for motor only models (Motor-Models: LR (WRS, P=0.001), UD (WRS, P=0.005), and 2D (WRS, P=0.069)). Line plots (**A**): Shaded area represents +/- 1 standard error of the mean (S.E.M). Violin plots (**B**): Shaded area represents kernel density estimate of the data, white circles represent the median, and grey bars represent the interquartile range.

The different decoding methods differed significantly in their performance; averaging across all sessions for each subject, a main effect of classification type was found (Figure 2B, F_1,60_= 21.69, P<0.00001; Supplementary Table S1). Post-hoc tests revealed that the use of all electrodes in CovNet models significantly improved classification accuracy compared to online performance across all tasks examined (LR: +11.79%, P=0.00002; UD: +11.47%, P=0.00002; 2D: +5.35%, P=0.006; Supplementary Table S2). CovNet models using only the motor electrodes displayed significant improvements in the 1D (LR and UD), but not the 2D, tasks when compared to the online classification accuracy (LR: +7.47%, P=0.001; UD: +7.19%, P=0.005; 2D: +3.52%, P=0.069; Supplementary Table S3). However, CovNet models using all electrodes did not yield significantly better classification accuracy than CovNet models using only motor electrodes (LR: +4.32%, P=0.21; UD: +4.28%, P=0.15; 2D: +1.83%, P=0.33; Supplementary Table S4).

When examining the relationship between online BCI performance and CovNet classification accuracy, it was found that the use of CovNet Models preferentially benefitted participants that struggled with online control. A significant positive correlation was found between online accuracy and CovNet classification, indicating that if participants produced better linearly classifiable signals, higher classification accuracy could be expected from the CovNet models (Figure 3A; Motor-Models: r_62_=-0.84, P<0.00001; All-Models: r_62_=-0.73, P<0.00001). A significant negative correlation was found between online BCI performance and the difference between CovNet classification accuracy and online BCI performance for both models (Figure 3B; Motor-Models: r_62_=-0.35, P=0.005; All-Models: r_62_=-0.54, P<0.00001). This suggests that, on average, the worse a participant performed during the standard online SMR-based BCI, the more benefit could be expected from using CovNet classification. Finally, the improvement in classification accuracy when using all electrodes to train CovNet models, vs just motor electrodes, displayed a significant negative correlation with online performance, indicating that even though these participants may not display the expected motor correlates of motor imagery, deep learning methods were able to find valuable information outside the motor cortex to aid in classification (Figure 3C; r_62_=-0.53, P<0.00001).

**Figure 3.**
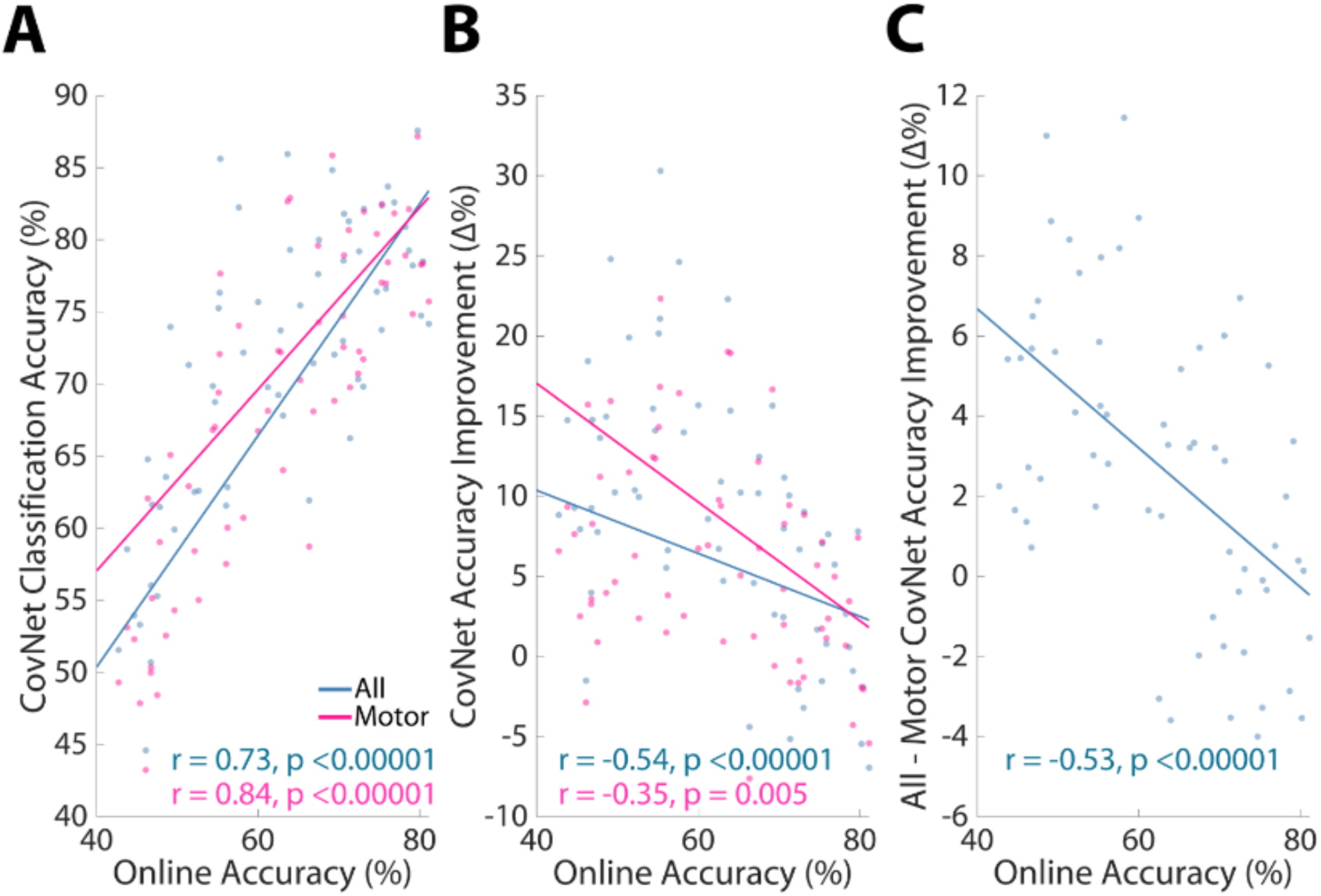
CovNet Classification Preferentially Benefits Inefficient Participants. **A**, A significant positive correlation was found between online BCI accuracy and CovNet classification accuracy for both All-Models (Pearson correlation, P < 0.00001) and Motor-Models (Pearson correlation, P < 0.00001). **B**, A significant negative correlation was found between online BCI accuracy and the difference between CovNet classification accuracy and online BCI performance for both All-Models (Pearson correlation, P < 0.00001) and Motor-Models (Pearson correlation, P = 0.005), suggesting those with worse online BCI accuracy improved more through the use of deep learning methods. **C**, Another significant negative correlation was found between online BCI performance and the improvement in CovNet classification accuracy when using All-Models compared to Motor-Models (Pearson correlation, P < 0.00001), indicating that those with worse online BCI accuracy improved more through the use of deep learning models with full scalp EEG coverage.

### 3.2. Using Estimated Class Probability as a Control Signal Highlights the Merit of High Density EEG

One important aspect of SMR-BCI is its continuous control; therefore, attempts were made to emulate online control with the output of the CovNet models. In BCI2000 (and standard cursor control tasks), the magnitude of cursor movement is determined by the strength of the chosen control signal (originally as normalized differences in alpha power over the motor cortex; hence larger differences in alpha power would lead to larger cursor movements). We simulated continuous control using the estimated class probability output by the CovNet models (i.e., one prediction for each 40ms interval), such that when the CovNet model was more confident in its decisions, the cursor could be moved further in a simulated environment.

In this simulation of continuous control, the CovNet models using all electrodes outperformed the models using only motor electrodes. Averaging across all 40ms intervals, the probability of correct class membership (e.g., all “left” probabilities averaged across all windows in “left” trials) was found to be significantly greater in the All-Models output compared to the Motor-Models (Figure 4A; F_1,60_=20.27, P=0.00001; LR: +0.05, P=0.009; UD: +0.07, P=0.003; 2D: +0.03, P=0.02; Supplementary Table S5,S6). Thus, on average, and for each increment of 40ms, a simulated cursor could be moved farther in the correct direction when including information across all available electrodes.

**Figure 4.**
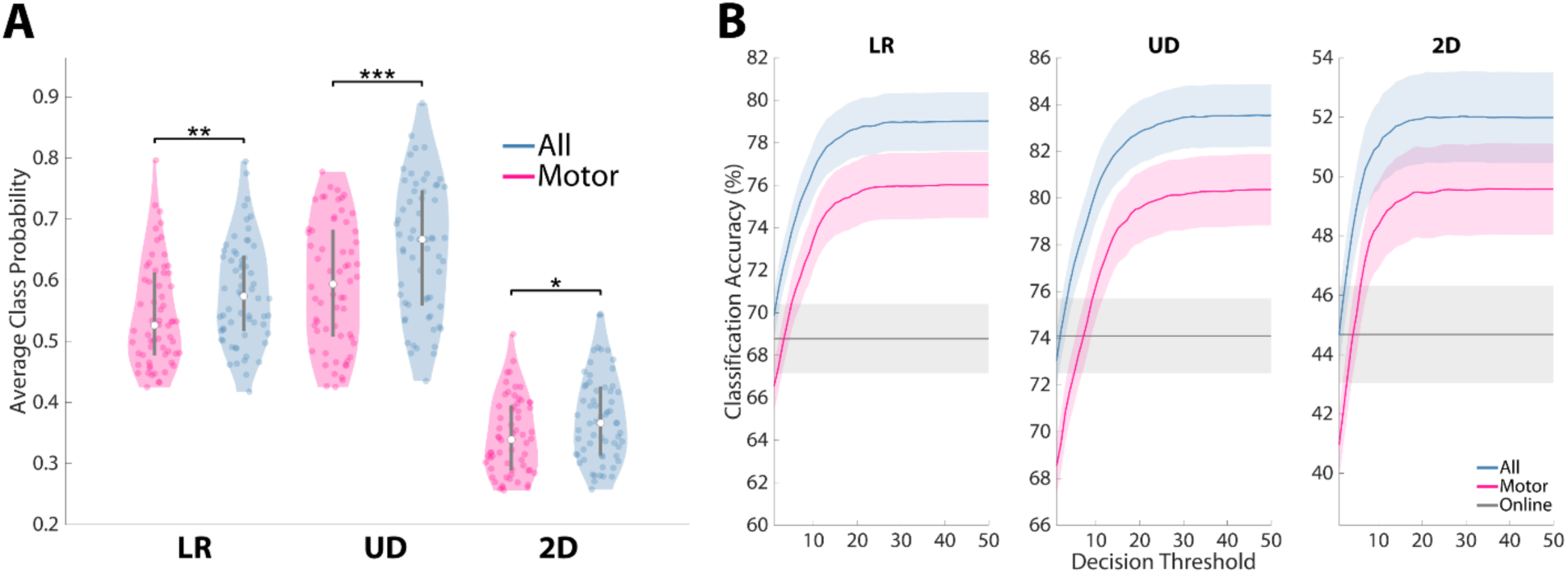
CovNet Models With Full Scalp Coverage Display a Favorable Theoretical Control Signal. Using a sliding window approach, the output of CovNet models (the predicted probability of class membership) can be used as a control signal for continuous BCI. **A**, When averaging across all moving windows, CovNet models with full scalp coverage display a higher probability of correct class membership across all tasks examined when compared to motor only models (LR (WRS, P=0.009), UD (WRS, P=0.003), and 2D (WRS, P=0.02)). **B**, A receiver operating characteristic curve was made by varying a threshold of the summed probability signal for classification (i.e. the probability of class membership of each window is summed from the beginning of a trial, then a classification occurs when this summed signal crosses the chosen threshold). When comparing the area under these curves, a main effect was found for model type (RAOV, P = 0.015), however significant differences were not observed in the post-hoc tests (LR (WRS, P=0.1), UD (WRS, P=0.052), and 2D (WRS, P=0.18)). Violin plots (**A**): Shaded area represents kernel density estimate of the data, white circles represent the median, and grey bars represent the interquartile range. Line plots (**B**): Shaded area represents +/- 1 standard error of the mean (S.E.M). The grey horizontal band represents the average online classification accuracy.

Since the models using all electrodes were “more confident” in their decisions, and this effect was compounded every 40ms, we believed a cursor could be moved faster in a simulated environment without substantially sacrificing the resulting classification accuracy. A simulated cursor control environment was created to examine the effects of deep learning classification on continuous BCI control where a virtual cursor was moved in a 2D plane similar to the feedback provided to participants during online control. For each EEG data window, a simulated cursor was moved in the direction of, and in direct proportion to, the highest class probability output by the CovNet models. In the same manner as online BCI control, left and right (as well as up and down) decisions were given opposite signs such that if two consecutive decisions were 0.65 probability Right, then 0.65 probability Left, these decisions would cancel (however, unlike online control, motion was restricted to a single axis per time window). If this summed signal crossed a threshold (varied across simulations) in one direction, a classification was made and the remaining trial data was discarded. If this threshold was not reached by the end of the trial data, the target closest to the simulated position was chosen as the model’s prediction.

Figure 4B displays a receiver-operating-characteristic curve plotting average classification accuracy against the classification threshold used for the summed probability signal. The x-axis of figure 4B can be thought of as “window width” in the simulated environment with lower values representing smaller distances between the targets (and fewer correct/incorrect decisions leading to target selection) and larger values representing wider windows and larger distances between targets (where more correct/incorrect decisions are needed for the cursor to contact a target). This is analogous to varying cursor speed in online scenarios where the screen width would be fixed, and hence lower values in the x-axis of Figure 4B can be thought of as representing faster cursors and larger values representing slower cursors. When comparing the area under the curve for these models, a main effect of model type was found (F_1,60_=5.96, P=0.015; Supplementary Table S7) however, none of the individual post-hoc tests reached significance (LR: +0.92, P=0.10; UD: +1.15, P=0.052; 2D: +0.73, P=0.18; Supplementary Table S8).

By emulating the tuning of cursor movement parameters in proportion to the CovNets’ output probability by varying the summed probability threshold needed for a classification, we attempted to find the point of optimal speed-accuracy tradeoff. We hypothesized that even though the increases in classification accuracy, probability of correct class membership, and AUC provided by the use of all electrodes were modest, these benefits could be traded for substantial improvements in speed of operation. First, in order to determine the operation point of the most accurate system using only motor electrodes, the decision threshold that maximized the Motor-Models’ classification accuracy was first identified (with the maximum found at or before the plateau where longer trials had no effect on average classification accuracy). At this threshold, significant reductions, compared to online control, in average trial length (LR: −0.56s, P<0.00001; UD: −0.38s, P=0.0004; 2D: −0.24s, P=0.005; Supplementary Table S9) and increases in ITR (LR: +2.14bits/min, P=0.0005; UD: +3.23bits/min, P=0.002; 2D: +1.65bits/min, P=0.016; Supplementary Table S10) were observed.

Then, in order to leverage the benefit provided by the additional information contained in full scalp EEG, the decision threshold which produced an equivalent classification accuracy for the All-Models’ output was chosen, therefore sacrificing a few percentage points worth of accuracy for a theoretical increase in speed of operation. In this way, we believed we could demonstrate a theoretical BCI system which optimally balances accuracy (i.e., the maximum achievable accuracy provided by motor only CovNet models) and speed of operation.

A significant interaction was observed between model type and BCI task when examining average trial length (Figure 5A; F_1,60_=17.18, P<0.00001; Supplementary Table S11). The average trial length of the All-Models’ output in a simulated environment was found to be significantly lower than both online control (LR: −2.51s, P<0.00001; UD: −2.07s, P<0.00001; 2D: −2.05s, P<0.00001; Supplementary Table S12) and Motor-Models (LR: −1.97s, P<0.00001; UD: −1.68s, P<0.00001; 2D: −1.80s, P<0.00001; Supplementary Table S13). ITR was then calculated with the classification accuracies and average trial lengths derived when using the CovNet Models’ summed probability thresholds, and a main effect for model type was found (Figure 5B; F_1,60_=94.42, P<0.00001; Supplementary Table S14). The ITR realized by tuning the All-Models’ probability output was significantly greater than the ITR of online control. (LR: +9.44bits/min, P<0.00001; UD: +12.49bits/min, P<0.00001; 2D: +7.57bits/min, P<0.00001; Supplementary Table S15). More importantly, this rate was found to be twice that of the Motor-Models (LR: +7.30bits/min, P<0.00001; UD: +9.27bits/min, P<0.00001; 2D: +5.92bits/min, P=0.00001; Supplementary Table S16).

**Figure 5.**
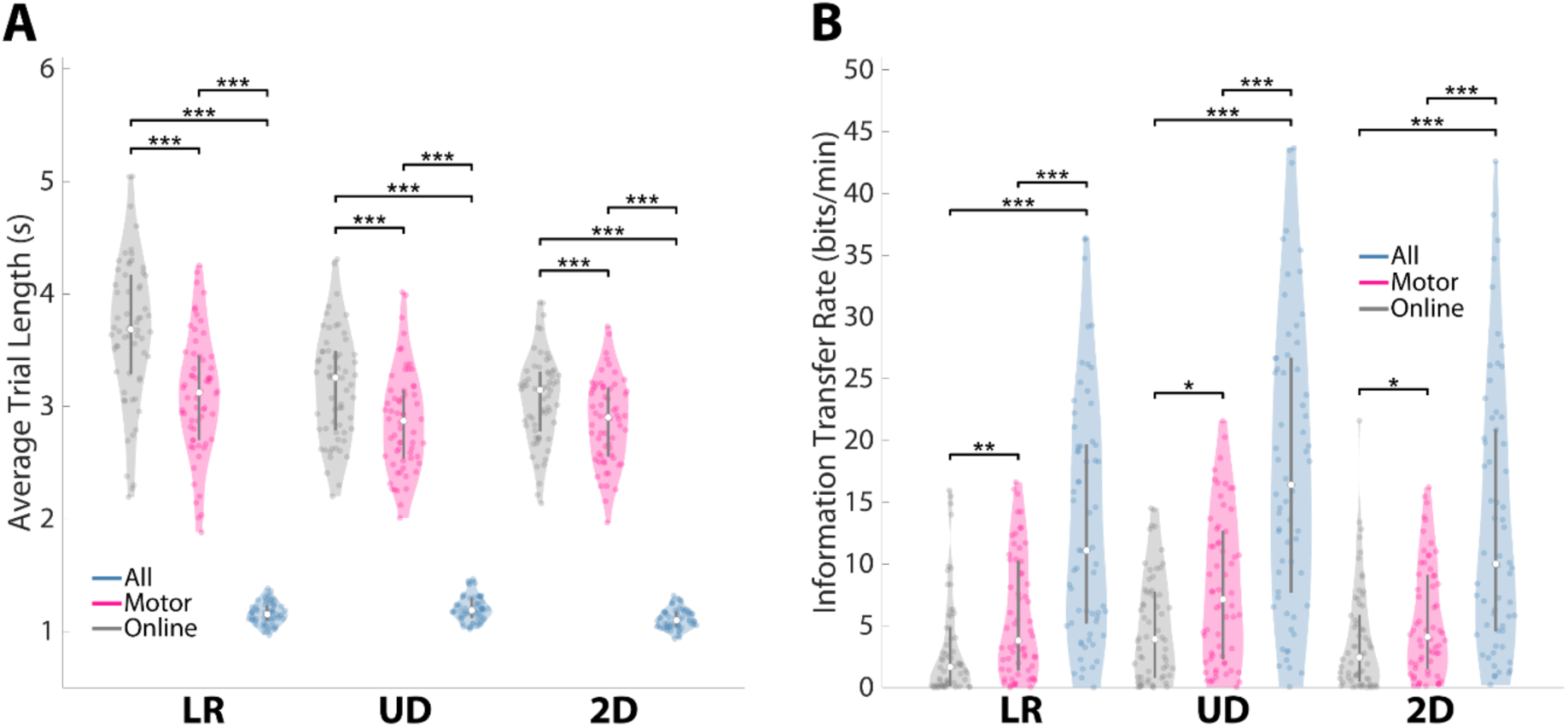
CovNet Models Significantly Increase Information Transfer Rate By Reducing Average Trial Length. Using a summed confidence threshold (Figure 3) for trial classification that maximized classification accuracy resulting from Motor-Model prediction and maintained this classification accuracy for All-Model prediction, CovNet classification was found to **A**, significantly reduce the average trial length (Vs. Motor-Models: LR (WRS, P<0.00001), UD (WRS, P=0.0004), and 2D (WRS, P=0.005), Vs. All-Models: LR (WRS, P<0.00001), UD (WRS, P<0.00001), and 2D (WRS, P<0.00001)) and **B**, significantly increase ITR (Vs. Motor-Models: LR (WRS, P=0.0005), UD (WRS, P=0.002), and 2D (WRS, P=0.016), Vs. All-Models: LR (WRS, P<0.00001), UD (WRS, P<0.00001), and 2D (WRS, P<0.00001)) when compared to online control in a simulated environment. The use of All-Models compared to Motor-Models displayed the same trends for average trail length and ITR (all, P<= 0.00001). Violin plots (**A**,**B**): Shaded area represents kernel density estimate of the data, white circles represent the median, and grey bars represent the interquartile range.

### 3.3. Feature Importance Maps Suggest CovNets Utilize Motor and Non-Motor Biomarkers for SMR-BCI Classification

The feature maps calculated with the input-perturbation network-prediction correlation approach (See section 2.4) revealed that both motor and non-motor biomarkers were utilized in the CovNet Models’ predictions (Figure 6). Cluster based permutation tests were used to compare the feature importance maps for each target against the feature importance maps for the other targets (while holding the frequency band fixed) in order to discover statistical regularities in the derived CovNet biomarkers for each class condition. The bands examined were (upper) alpha (10-14Hz), Beta (16-28Hz), and Gamma (28-58Hz). In total, 26 clusters were identified and all clusters displayed a significance level lower than the Bonferroni threshold for multiple comparisons (P<0.0042) (see Supplementary Figure S2, Supplementary table S17). Right hand motor imagery was associated with patterns of contralateral motor and ipsilateral occipital desynchronization in the alpha and beta bands with a unique ipsilateral fronto-tempolar gamma response. Left hand motor imagery displayed the same pattern mirrored over the midline. Motor imagery of both hands correlated with motor and occipital desynchronization across all bands as well as with frontal beta synchronization. Resting was associated with alpha and beta synchronization over the motor cortex as well as frontal beta and frontal/temporal gamma desynchronization.

**Figure 6.**
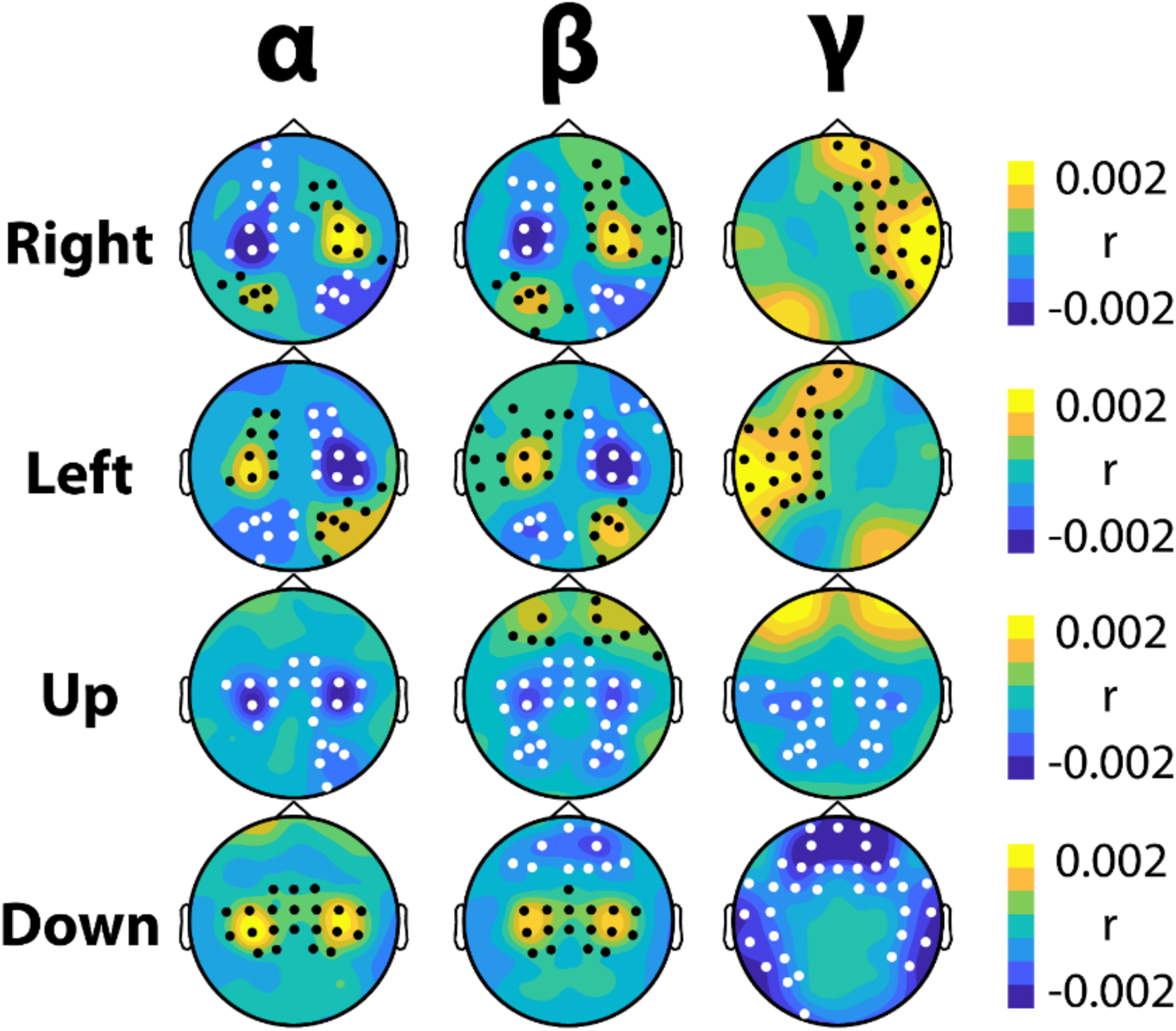
Feature Importance Maps Display Multiple Biomarkers Used in Continuous BCI Control. Using the input-perturbation network-prediction correlation method, feature importance maps were created for each motor imagery strategy. Warm colors occur when increases in power over these areas increase the classification accuracy, while cool colors occur when decreases in power increase the classification accuracy (or vice-versa). Black dots represent significant clusters of positive correlation and white dots represent significant clusters of negative correlation. All clusters had a significance level lower than the Bonferroni threshold for multiple comparisons (P<0.0042).

Following the finding of a significant negative relation between improvement in classification accuracy and online control (Figure 3B), it was hypothesized that the participants who displayed the greatest improvements in classification accuracy would have a stronger dependence on features external to the motor area, specifically occipital areas during horizontal (Left/Right) BCI control. Thus, an exploratory analysis was conducted by splitting participants into two groups: those who improved most through the use of CovNet classification, and those with the least improvement. Cluster based permutation tests comparing the feature importance maps of these two groups within the alpha, beta, and gamma band revealed that there was a significantly stronger negative relation between ipsilateral occipital beta power and network prediction in the most improved group (Figure 7A; Right Occipital Beta—cluster stat (maxsum(*t*_60_))=-7.44, P=0.025; Left Occipital Beta—cluster stat (maxsum(*t*_60_))=-8.47, P=0.018; not corrected for multiple comparisons). This finding suggests that ipsilateral occipital beta power was more important in CovNet classifications in the group that benefitted most through the use of deep learning. To determine how this feature differed between the two groups, EEG data was time-locked to moments of the greatest cursor movement. When averaging beta power over the identified clusters, it was indeed found that the participants that most benefitted from CovNet classification displayed greater reductions in ipsilateral beta power roughly 300ms after the moments of greatest cursor movement (Figure 7B; R Occipital Beta—cluster stat (maxsum(*t*_60_))=-46.93, P=0.003; L Occipital Beta—cluster stat (maxsum(*t*_60_))=-17.92, P=0.017).

**Figure 7.**
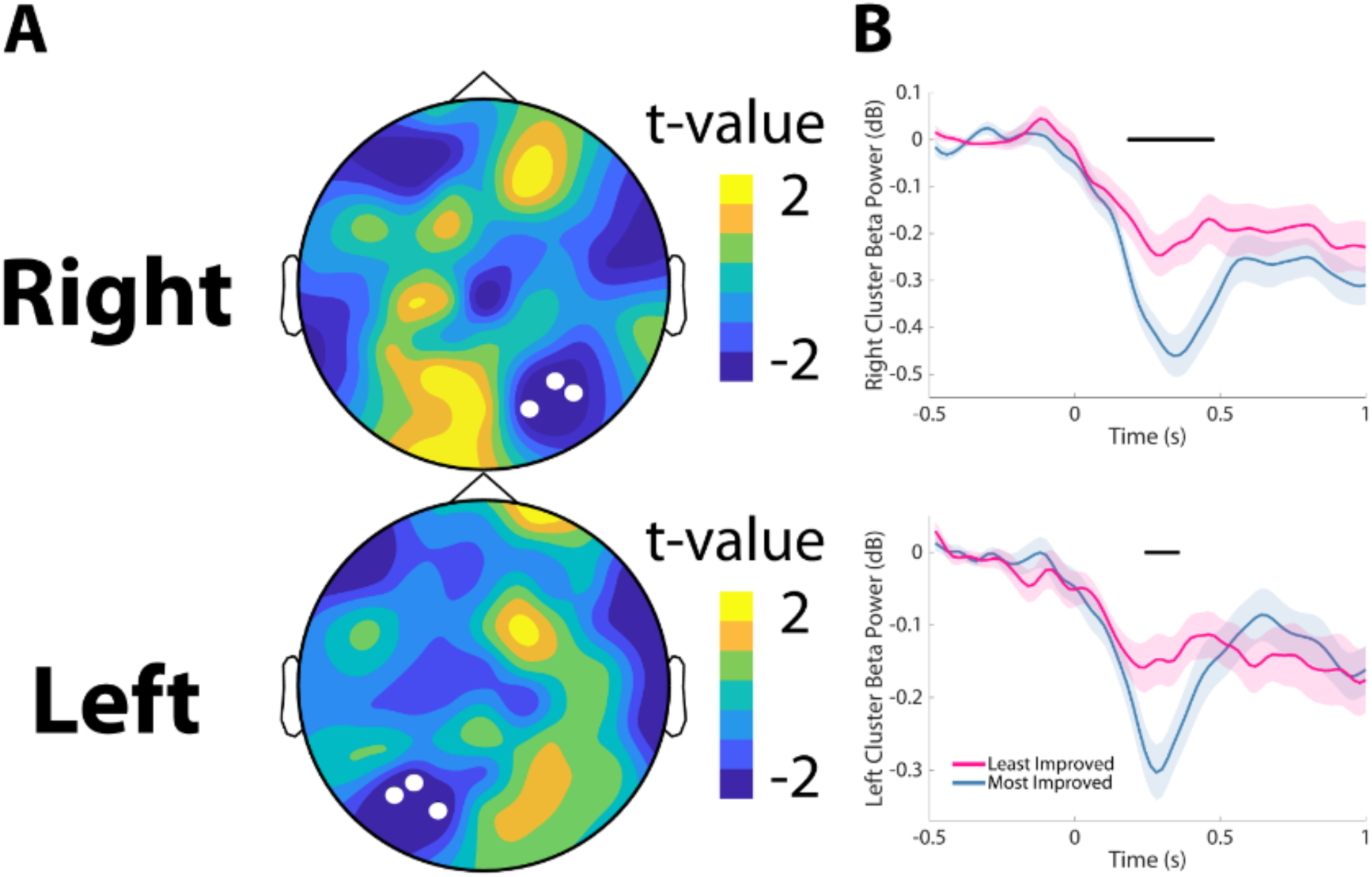
Subjects Benefitting from CovNet Classification Display More Distinct Visual Features. **A**, Ipsilateral occipital beta desynchronization was found to correlate more strongly with classification accuracy in participants that most improved through the use of CovNet classification when comparing feature importance maps between these populations (R: Cluster based permutation test (CBPT), P=0.025; L: CBPT, P=0.018). **B**, When compared to the population of participants that improved least, the most improved participants display greater ipsilateral occipital beta desynchronization in response to cursor movement (R: CBPT, P=0.003; L: CBPT, P=0.017).

## 4. Discussion

In this work, we have demonstrated that deep learning methods are able to significantly improve classification accuracy in SMR based BCI tasks. This improvement was found to disproportionately benefit the participants who struggled the most with BCI control, suggesting the use of CovNet classification in BCI could potentially address the problem of BCI inefficiency. The use of high density EEG was shown to significantly improve the theoretical speed of operation and ITR (without sacrificing classification accuracy) when compared to models utilizing information derived from traditional “motor only” electrode montages. Finally, novel combinations of BCI biomarkers were identified in addition to standard sensorimotor rhythms, such as occipital features and frontal gamma power, indicating deep learning models may derive useful information from outside the motor cortex when classifying EEG data collected during online SMR MI BCI control with continuous feedback.

Recent research demonstrating improved classification accuracy through the use of deep learning methods was extended here in a large and longitudinal BCI dataset. Most of the recent work in applying neural networks to SMR based BCI has focused on the development of various architectures and algorithms while testing them on standard benchmark, or in-house, datasets (Lawhern et al., 2018; Lu et al., 2017; Sakhavi et al., 2018; Schirrmeister et al., 2017; Tangermann et al., 2012; Wang et al., 2018), though some have recently begun to validate these methods on their own datasets in online scenarios (Jeong et al., 2020; Tayeb et al., 2019). Here, we have shown that deep learning with convolutional neural networks can significantly increase classification accuracy on an independent, large, and longitudinal dataset, thus validating the generalizability of these methods. We additionally demonstrate that these methods are able to produce high classification accuracy in scenarios suitable for continuous SMR based control, as opposed to prior work that produced one classification per uniform 2 or 4s trials (Lawhern et al., 2018; Lu et al., 2017; Sakhavi et al., 2018; Schirrmeister et al., 2017; Wang et al., 2018).

The inability of a significant proportion of the population to control standard BCIs, even after extensive training, has historically limited the clinical adoption of BCI technology (Leeb et al., 2013). However the use of CovNet models in BCI systems could potentially remedy this situation as the improvement in classification accuracy through the use of deep learning methods was discovered to be greatest in this challenged population. This finding may lend credence to the future clinical viability of BCIs as BCI decoding technology develops (Guger et al., 2017). Those with poorer initial BCI performance additionally displayed a greater benefit from the inclusion of extra-motor electrodes in this offline classification study, suggesting high-density EEG may be more important for end-users. Most prior work has focused on classification which uses electrodes only over the motor cortex, with some claiming the inclusion of all electrodes for BCI classification was found to lead to worse accuracies than those over the motor cortex alone (Schirrmeister et al., 2017), therefore future work is needed to reconcile these discrepant findings.

BCI control makes use of a distributed network of interacting brain structures that participate in motor imagery, maintenance of task demands, attention, awareness of feedback, and reward processing (Sitaram et al., 2017). As predicted, characteristic motor cortex dependent patterns of motor imagery within the alpha band, which additionally extended into the beta band, were found in the feature importance maps, demonstrating CovNets are able to find subject specific sensorimotor rhythms.

The alpha rhythm, and indeed performance in sensorimotor rhythm BCIs, has been shown to be related to gamma band activity, especially in the frontal regions (Ahn et al., 2013a; Grosse-Wentrup et al., 2011; Grosse-Wentrup and Schölkopf, 2014, 2012). This frontal gamma signal was found to play an important role in CovNet models, and, of particular note, its absence was a prominent biomarker of the rest condition where motor imagery was not performed. However, these maps additionally suggest that CovNet models utilize non-motor related features in their decisions, such as occipital signals.

Parietal/occipital attention signals have been used to develop various BCI techniques such as overt and covert spatial attention (Bahramisharif et al., 2010; Meng et al., 2018; Tonin et al., 2013; Treder et al., 2011; van Gerven and Jensen, 2009). While these studies focus on synchronization of the alpha rhythm subsequent to shifting attention, their results also indicate there is a pattern of mid-frequency desynchronization during the initial shifting period (Treder et al., 2011; van Gerven and Jensen, 2009). Ipsilateral occipital alpha and beta desynchronization were found to be significantly correlated with CovNet model predictions, which alludes to a pattern of orienting to the target while covertly attending to the cursor (opposite that of covert attention paradigms of orienting to the center of the screen while covertly attending to the periphery) (Bahramisharif et al., 2010; van Gerven and Jensen, 2009). This ipsilateral occipital beta signal was shown to play a more prominent role during LR control in models trained on participants that benefitted most from CovNet classification. Moreover, these participants were found to display a stronger beta desynchronization response following cursor movement. While the group-averaged feature importance maps include all of the aforementioned biomarkers, individual models may rely on a subset of these biomarkers, or even unique features that do not appear when examining BCI data from the group level. One advantage of CovNet classification of BCI signals is the ability of these models to detect reliable biomarkers at the individual, rather than the group, level. It is this specificity which might explain why subjects who did not fit the “standard” BCI model had the greatest improvements in classification accuracy. As evidenced here, future probing of these deep-learning models might serve to discover relationships between subject populations and biomarkers leading to novel neuroscientific findings, such as when are these occipital and frontal gamma features are in operation, which subjects display them most prominently, and whether they can be trained or enhanced.

The significant increases in classification accuracy suggest future BCIs using CovNet classification could be controlled with finer precision, however the robustness of the control signal—as opposed to its precision—may lead to greater practical advantages. Since more information was summarized in the control signal when using high density EEG, we believed we could increase the speed of cursor movement in a simulated environment without sacrificing classification accuracy. We indeed found that significant reductions in average trial length and increases in ITR could be achieved by tuning the CovNet models’ confidence estimations in a simulated environment. Further, when the slight advantage provided by All-Models was traded for increases in speed, high-density EEG was found to double the ITR without any loss of classification accuracy when compared to a motor-only montage. By increasing ITR through the use of deep learning classification, future online BCI systems may be able to issue more commands in shorter time. Increases in ITR through the use of CovNet classification have been documented in other, non-motor imagery based BCI paradigms, which suggests EEG signals may contain more information than generally assumed (Nagel and Spüler, 2019).

Challenges still exist when translating CovNet BCIs from offline classification analyses to online systems. One limitation of this study is that, while we believe it represents one of the most complex classification studies conducted, these analyses were still performed offline. In online experiments, training data may not be available beforehand, or if it was collected prior to BCI use, the training of models within an experimental session would take a prohibitive amount of time. Future work will be needed to assess how well models trained in one session transfer to future sessions (Fahimi et al., 2019). Additionally, neural networks often require long training times themselves, so there is a need to devise systems that are able to adapt in an online fashion (Schwemmer et al., 2018). For example, variable electrode placement may compromise the accuracy of models trained in previous sessions, which is a hindrance not present in static, invasive systems (Schwemmer et al., 2018). Recently, the feasibility of online BCI systems using deep learning classification has been demonstrated (Jeong et al., 2020; Tayeb et al., 2019), however more work is needed to create noninvasive deep learning BCIs that provide continuous control. BCI is a skill that the user and system acquire together, and it is currently unclear how neural networks will adapt to changes in a user’s control strategy as it develops (Perdikis et al., 2018; Vansteensel et al., 2016).

## 5. Conclusions

In conclusion, we have demonstrated that deep-learning based classification and high density EEG are important tools for increasing BCI efficiency. CovNet models are shown to significantly increase the classification accuracy of motor imagery on a large BCI dataset. This improvement is found to have the greatest benefit in the most challenged subset of the population, namely those that struggle with standard BCI control, suggesting deep learning in BCI systems may have the greatest impact on end-users in the locked-in state (De Massari et al., 2013; Grosse-Wentrup and Schölkopf, 2014; Kübler and Birbaumer, 2008). BCI inefficient participants were again discovered to disproportionately benefit from the use of full scalp EEG coverage. Further, leveraging the output of these models in a simulated cursor control environment demonstrated that the additional information provided by high-density EEG could substantially increase the ITR without any loss in classification accuracy (when compared to motor-only electrode montages). This benefit may be in-part due to the variety of reliable correlates of BCI control, in addition to the sensorimotor rhythm (such as an occipital beta signal in response to cursor movement) that may be learned. These findings additionally suggest that valuable information may remain latent within EEG recordings (Nagel and Spüler, 2019). Future work developing BCI systems with online classification using CovNet models may lead to the widespread clinical adoption of this technology and aid those suffering with paralysis by restoring their agency and access to the world.

## Acknowledgements

We would like to thank Dr. Mary Jo Krietzer and Dr. Christopher C. Cline for their assistance in subject recruiting and data collection, and Rui Sun for useful discussions on data analysis.

## Funding

This work was supported in part by the National Institutes of Health (grants AT009263, MH114233, EB021027, NS096761, and EB008389).

## Author contributions

B.H., S.E., and J.R.S. designed the experiments. J.R.S. collected the data. J.R.S. analyzed the data. D.S. provided methodological guidance. J.R.S., S.E., D.S., and B.H. wrote the paper. All authors discussed the results and contributed to editing the manuscript.

## Competing interests

The authors declare no competing interests in the work reported here.

## Footnotes

1 Abbreviations: BCI—Brain computer interface, SMR—Sensorimotor rhythm, ITR—Information transfer rate

## Notes

### Competing Interest Statement

The authors have declared no competing interest.

